# Modeling Microbiome Modulation of Tumor Metabolic Networks to Predict Synergistic Therapies

**DOI:** 10.64898/2026.02.25.707963

**Authors:** Annie J Badenoch, Zeyang Pang, Carolina H. Chung, Aaron Robida, Bretton Badenoch, Ritish Natesan, Layth Kakish, Jiahe Li, Sriram Chandrasekaran

## Abstract

Differences in microbiome composition profoundly influence drug response, yet methods to model the metabolic interplay between tumors, microbes, and therapeutics remain limited. We present a generalizable framework combining machine-learning and genome-scale metabolic modeling to prioritize combination therapies for colorectal cancer (CRC) in the presence of *Fusobacterium nucleatum* (*Fn*) and other pathogenic, probiotic, and commensal microbes. Trained on 6,514 drug combinations in microbe-free CRC cell lines, the model predicted synergistic combinations in both microbe-free and microbe-associated contexts and generalized to immunotherapy-associated conditions. Predictions were validated using an asymmetric co-culture system that mimics the colon’s normoxic–anaerobic gradient, confirming synergistic combinations in HCT116 cells with *Fn*, including drugs not typically used in CRC therapy. Mechanistic analysis and targeted pharmacological perturbations revealed phospho-inositol metabolism and cysteine transport as key determinants of *Fn*-dependent drug synergy. Together, this work introduces a scalable, microbiome-aware framework to enable discovery of context-specific combination therapies.

## Introduction

There has been growing interest in understanding the impact which microbes have on cancer metabolism and subsequently cancer progression and treatment outcomes.^1^ Interpatient variability in treatment response is due in part to individual variation in the microbiome in the tumor microenvironment (TME).^2^ Several microbes have been implicated in cancer progression and drug resistance, with *Fusobacterium nucleatum* (*F. nucleatum, Fn*) having particularly strong evidence in Colorectal Cancer (CRC).^3–5^ In the United States, CRC affects 4% of people, with 15-20% of patients experiencing chemotherapeutic drug resistance, and 30-40% facing cancer recurrence.^6^ The high rates of chemotherapy resistance, coupled with the rising incidence of CRC in patients under 55, underscore the urgent need for new therapeutic strategies.^6–8^ Although multi-agent regimens such as the combination of folinic acid (leucovorin), 5-fluorouracil, and oxaliplatin (FOLFOX) have improved outcomes, 50–70% of metastatic CRC patients remain non-responsive.^9,10^ Interpatient variability in treatment response is due in part to individual variation in the microbiome in the tumor microenvironment.^2,11–14^ These complexities present a major hurdle in treating CRC and require more dynamic, patient-specific approaches.

To address this challenge, we investigated how therapeutic agents and tumor-associated microbes modulate CRC metabolism. CRC cells exhibit elevated glycolytic activity and dynamically reprogram their metabolism based on oncogenic mutations, nutrient availability, and environmental pressures.^1^ For example, under conditions of glutamine depletion, CRC cells can rewire their metabolic programs to favor asparagine biosynthesis.^15–17^ Many first-line CRC therapies directly target metabolic pathways, including fluorouracil’s disruption of nucleotide biosynthesis.^18,19^ The strong link between CRC and metabolism serves as a basis for using metabolic perturbation to predict non-obvious therapeutic vulnerabilities.

The complexity of CRC treatment is further compounded by the metabolic influence of gut microbes, like *Fn*. Variations in microbiome composition have been linked to chemotherapy resistance and poor clinical outcomes.^20,21^ Although *Fn* is primarily an oral commensal and rarely detected in the healthy colon, multiple studies report its enrichment in CRC tumors.^3,4,22^ Additionally, *Fn* can metabolically modulate the cancer cells.^23,24^ *Fn*’s known relationship with colon cancer and metabolic impact makes it an ideal microbial species for investigating microbe-induced metabolic effects on drug and immunotherapy responses.

We hypothesized that these metabolic changes, which make CRC resistant to conventional chemotherapies, could be exploited to identify new, unintuitive drug combinations that counteract microbe-driven metabolic reprogramming. Modeling this three-way interaction among microbes, cancer cells, and therapeutic agents demands computational frameworks that can jointly capture metabolic constraints and drug-response patterns. To address the challenges of simulating the complex interplay of microbes, cancer cells, and drug response, here we applied a combination of machine learning (ML) and genome-scale metabolic modeling (GEMs). GEMs have been used to study microbiomes and microbe-host interactions, but have not been extended to predict drug responses or to identify synergistic drug combinations in the presence of specific microbes.^25–32^ Likewise, existing ML methods can predict drug-drug interactions but rarely incorporate microbial influences of microbe-drug combinations.^23,33–35^

Here we present a novel framework Onco-Microbiome-GEM-ML (OMG-ML), which integrates GEMs with ML to identify metabolic patterns associated with synergistic or antagonistic drug and microbe interactions. A key innovation of OMG-ML is its incorporation of microbe-induced metabolic modulation, allowing the model to account for the impacts of tumor-associated microbiome on drug response. In CRC, where both intrinsic drug resistance and microbe-driven modulation of treatment response are relevant, we hypothesized that OMG-ML can be used to identify drug combinations that are effective in tumor cells, both in standard conditions and in the presence of *Fn* and other microbes. OMG-ML is easily adaptable to other microbial species and can be continuously refined with emerging drug synergy data, enabling its application across diverse cancer types and biological contexts to discover microbiome-aware treatment combinations.

We first demonstrate that OMG-ML accurately predicts interactions between pairs of drugs and then extend it to simulate microbe-drug interactions and microbe-microbe-drug interactions. Notably, the model uncovered several highly synergistic drug combinations to treat CRC, including cabazitaxel paired with megestrol acetate. Significantly, it also identified drugs, such as methotrexate and fluorouracil, that exhibit enhanced efficacy in the presence of *Fn*. Moreover, the framework revealed agents like thalidomide and pralatrexate whose predicted effects vary depending on the microbial species present, highlighting the importance of incorporating microbe-specific metabolic modulation into therapeutic design. Collectively, these results establish OMG-ML as a powerful method for integrating metabolic modeling with ML to guide microbiome-aware treatment design and enable personalized drug combination discovery.

## Results

### Prediction of Chemotherapy Combinations

Flux balance analysis (FBA) of GEMs coupled with ML offers a mechanistic framework for identifying metabolic pathways underlying effective drug combinations. The GEMs enable simulation of pairwise and higher-order combinations of stressors, and ML identifies nonlinear flux patterns predictive of synergistic outcomes. In this approach, GEMs are constrained using transcriptomic data from individual drug treatments to generate treatment-specific metabolic states. These states are subsequently combined to represent the metabolic response to drug combinations, and the resulting flux-based features are used to train a random forest model on experimentally derived drug synergy scores.^36^

Our model, based on this technique, OMG-ML, was trained using transcriptomics data from 110 drug treatments on HCT116 and HT29 cells sourced from the library of integrated network-based cellular signatures (LINCS) and 6,514 chemotherapy combinations from SynergyxDB.^37,38^ HT29 cells are microsatellite stable, epithelial cells isolated from the colon of a female patient and have a less aggressive phenotype compared to HCT116 cells.^39^ HCT116 cells are microsatellite instable, epithelial cells isolated from the colon of a male patient and are often used to study CRC metastasis.^40^ Using these cell lines provides a diverse representation of CRC phenotypes to enhance the generalizability of the model across CRC subtypes. Individual drug treatment models were then combined to form composite feature vectors for each drug combination. An ensemble random forest was trained using these features and the known drug combination synergies from LINCS. A random forest was selected because of its superior performance compared to other ML models based on ten-fold cross-validation (synergy area under the curve (AUC) = 0.79, Antagonism AUC = 0.73, Pearson’s *r* = 0.54) (Supplementary Fig. 4f). The flux-based ML model also outperformed direct transcriptomics input (Supplementary Fig. 4a-e) and was highly significant compared to random permutations (*p*-value = 0.001) (Methods). Fig. 1 depicts a graphical overview of the development of OMG-ML. This framework enables prediction from human cellular metabolic flux profiles for previously unseen drug treatments and microbial co-culture conditions.

**Fig. 1.**
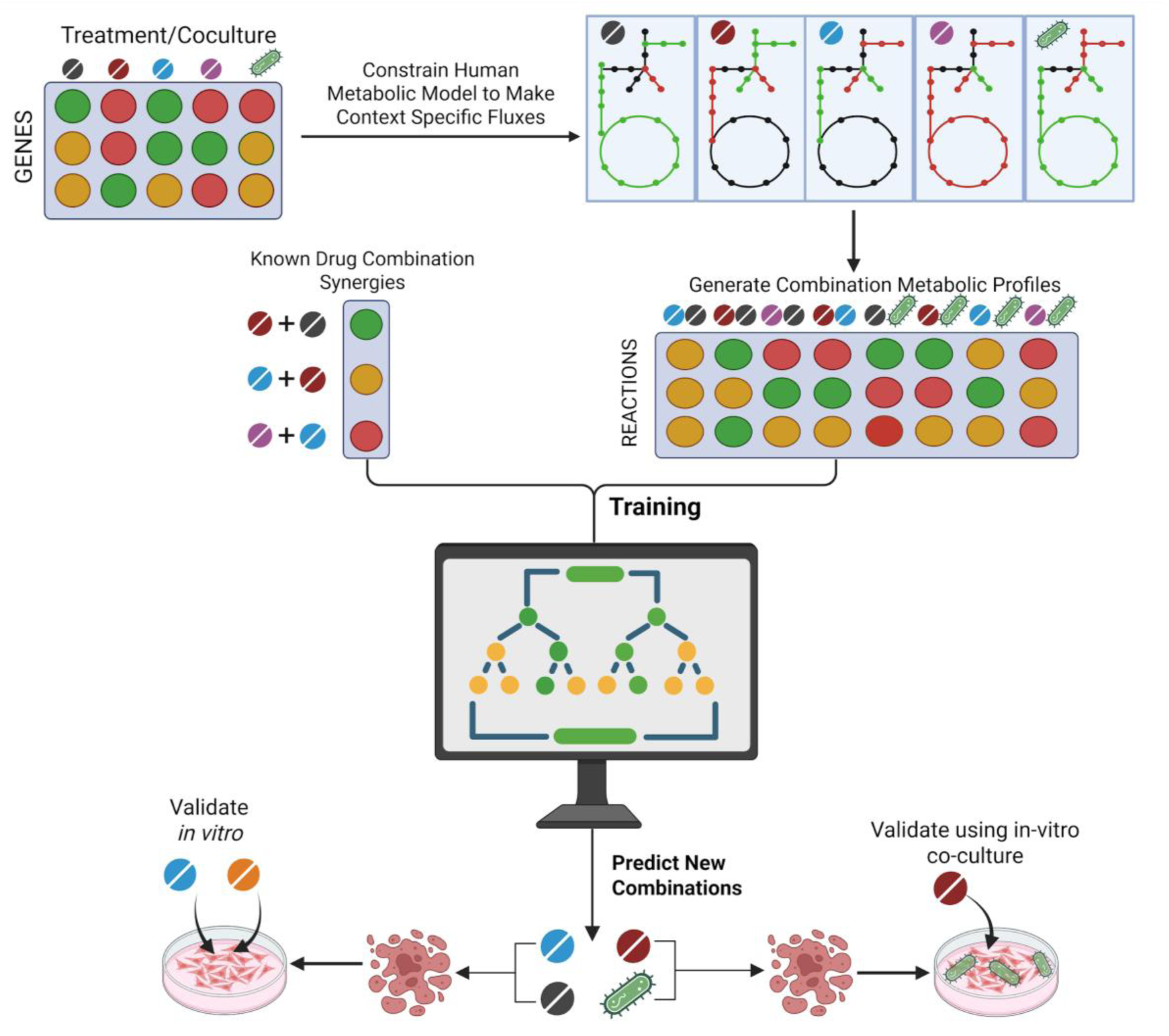
Overview of the OMG-ML workflow. Schematic illustrating model training and predictive output generation. Differentially expressed genes derived from drug treatment or microbial co-culture are used to compute metabolic flux profiles. These fluxes, together with known drug synergy measurements, are used to train the OMG-ML model. The trained OMG-ML framework is then applied to predict non-obvious drug combinations and identify drugs that are effective in the presence of specific microbes*. Figure created using BioRender.com*.

The trained model was used to predict 6,333 novel drug combination synergy scores (3,163 drug combinations on HCT116 cells and 3,170 on HT29 cells). Under the Loewe model, synergy is defined as negative values and indicates that the drug performed more strongly in the combination than alone. Synergistic combinations often included cabazitaxel (a taxane used to treat castration-resistant prostate cancer). The 15 most synergistic combinations on HCT116 cells were combinations including cabazitaxel (Supplementary Fig. 2). One of these synergistic combinations was cabazitaxel and megestrol, which was confirmed *in vitro* and showed strong synergistic effects (Fig. 2). Megestrol, or megestrol acetate, is a synthetic form of progesterone and has not been used as a colon cancer treatment, but has been used to increase patient appetite after chemotherapy.^41^ Of the predicted combinations, 11 were evaluated *in vitro* (Fig. 2a). These were combinations of cabazitaxel or megestrol with erlotinib, gemcitabine, metformin, methotrexate, and sorafenib. Predicted values and experimental values demonstrated Pearson’s correlation of 0.48, similar to the cross-validation evaluation, suggesting generalizable synergistic performance.

**Fig. 2.**
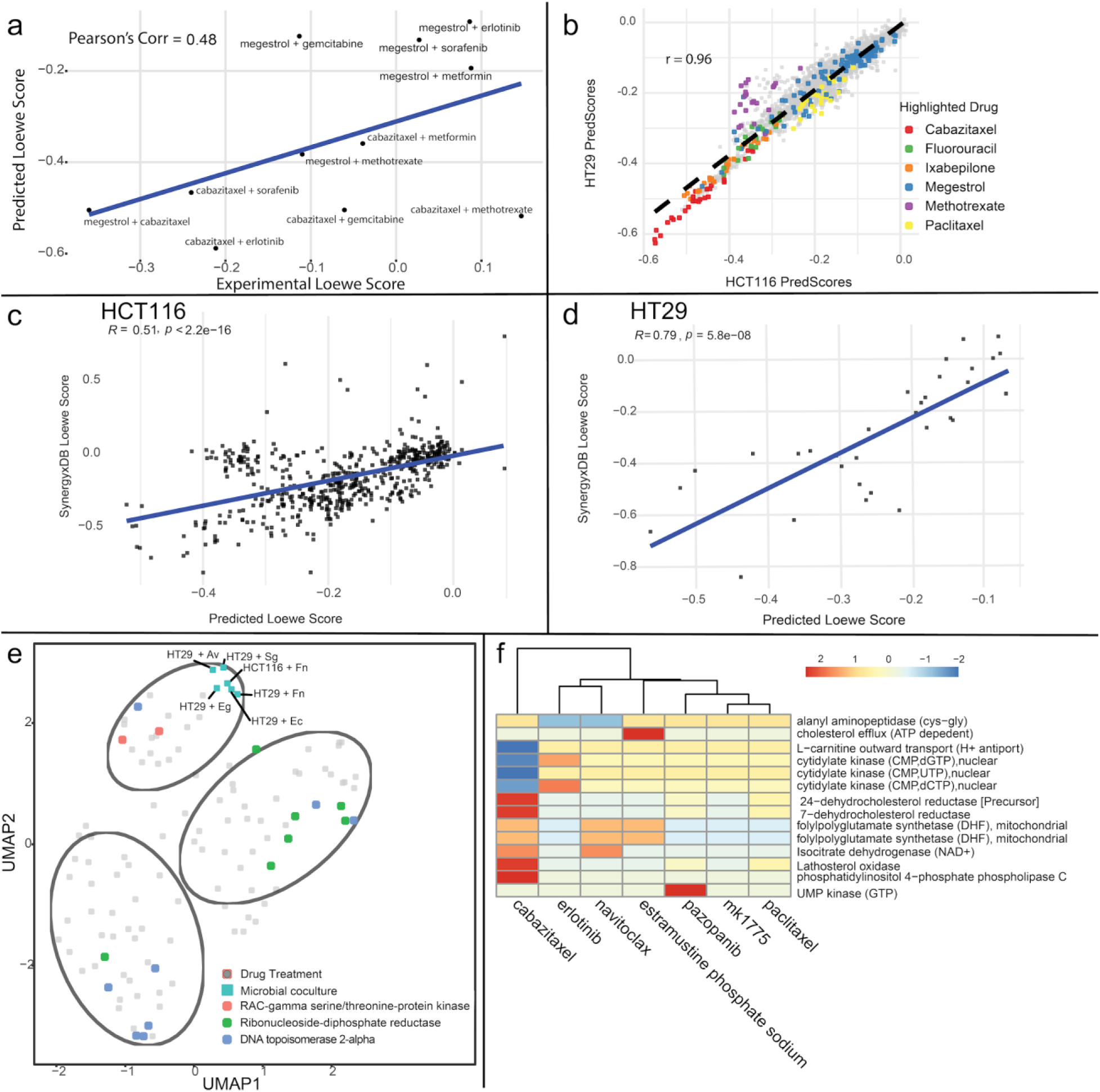
Predicted and validated drug synergy in HCT116 and HT29 cells with corresponding metabolic flux signatures. **(a)** Loewe scores from ML prediction for two drug combinations in the HCT116 cell line. Experimental scores from in vitro screening on the HCT116 cell line using SynergyFinder. **(b)** Predicted drug combinations in HCT116 and HT29 cells with drugs of interest highlighted. Colored points indicate the highlighted drug is one of the drugs in those two-drug combinations. Some of the cabazitaxel, megestrol, methotrexate, and fluorouracil combinations were validated in vitro. **(c)** Predicted Loewe scores from the model compared to known synergy scores from SynergyxDB for HCT116 cells (N = 554). Lower scores indicate stronger synergy. Significance of the correlation was found using the p-value for Pearson correlation. **(d)** Predicted Loewe scores from the model compared to known synergy scores from SynergyxDB for HT29 cells (N = 32). **(e)** Uniform manifold approximation and projection (UMAP) projection of treatment-associated flux changes. K-means clustering is overlaid as ellipses, and point colors denote the dominant enzymatic target characterizing each cluster. Flux responses induced by microbial treatments are highlighted using square-shaped markers. **(f)** Flux through enzymes targeted by each listed drug. Values were normalized using z-score transformation, calculated by subtracting the mean flux across all drugs in that pathway and dividing by the standard deviation. Reaction rows are clustered and not ordered based on importance.

To further test the model’s performance, predicted combination scores were compared to new scores from SynergyxDB. Since the initial development and training of the model, an additional 554 drug combinations have been reported for the HCT116 CRC cell line, along with 32 new combinations for the HT29 cell line. We applied our trained model to predict synergy scores for these newly added combinations and compared the predicted values to the experimentally measured synergy scores. The model’s predictions significantly correlated with these scores for both cell lines, with a Pearson’s *r* of 0.51 for HCT116 combinations and 0.79 for HT29 combinations (Fig. 2c-d).

The results are comparable to the performance of similar models. The CARAMeL model, which utilized similar feature engineering and flux-based approaches, also achieved a similar correlation on novel antibiotic combinations.^42^ Similar models that predict drug combination synergy for CRC such as by Tsiryouli *et al.*, which integrates multi-omics data and Consensus Molecular Subtype classifications achieved AUC values ranging from ∼0.5-0.78.^43^ Our model stands out from existing methods by its interpretability of pathways of interest, and its ability to account for the metabolic effects of tumors, chemotherapy, and microbiome.

Other combinations predicted by the model have support from literature. For example, MK1775 is a selective Wee1 kinase inhibitor that has been shown to improve the effects of fluorouracil in colon cancer cells.^42^ Our prediction of this combination aligns with this observation (predicted Loewe score = -0.31). These results provide support for the accuracy of our model’s predictions.

### Metabolic Insights on Drug Combination Treatments

As expected from prior studies, metabolic flux rewiring induced by individual drugs reflects their underlying mechanisms of action, with drugs with similar mechanisms of action clustering together (Fig. 2e).^36,44^

Because the ML model is trained using metabolic input, we next analyzed which reactions contribute most to the model’s predictive accuracy. By identifying the top-ranked features, we highlight specific metabolic reactions that are most strongly associated with effective drug combinations. Supplementary Fig. 1c depicts the normalized flux through the top 25 features for all drug treatments and classification of each drug as ‘synergistic’ or ‘antagonistic’. As shown in Supplementary Fig. 1c, using these features allows for separation between synergistic and antagonistic drugs.

These features can be used to understand the mechanism underlying why cabazitaxel was predicted to be more effective than paclitaxel (Fig. 2f). This comparison is important because paclitaxel and cabazitaxel are very mechanistically similar molecules, but elicited vastly different predictions.^45^ These top features are especially informative because cabazitaxel and paclitaxel exhibit very similar overall metabolic profiles (Euclidean distance = 4.2), compared to their average distances from other drugs (21.9 for cabazitaxel, 22.1 for paclitaxel). By focusing on flux differences in these top-ranked features, we can identify subtle but meaningful metabolic distinctions that may explain their divergent synergy predictions, despite their global similarity.

The five most synergistic drug combinations on HT29 cells were cabazitaxel combined with erlotinib, navitoclax, estramustine phosphate sodium, pazopanib, or MK1775. As shown in Fig. 2f, cabazitaxel uniquely decreased cytidylate kinase activity while increasing flux through several key reactions, including polyglutamate synthetase, phospholipase C, isocitrate dehydrogenase, dehydrocholesterol reductase, and lathosterol oxidase. Partner drugs showed variable effects across these pathways, generally exhibiting lower lathosterol oxidase and phospholipase C activity and higher cytidylate kinase flux relative to cabazitaxel alone.

### OMG-ML Predictions of Drugs in Combination with *Fn*

Given the growing recognition that colorectal cancer (CRC) is strongly influenced by the tumor-associated microbiome, *Fn* has emerged as a particularly relevant microbial factor due to its consistent enrichment in CRC and its reported roles in metabolic reprogramming and therapy response. To capture these microbiome-driven effects, the model was extended to incorporate the influence of microbes on cellular metabolism and drug response. This approach is based on the rationale that the metabolic effects of individual microbes resemble those of drug treatments, yet remain distinct, capturing microbe-specific perturbations (Fig. 2e). Furthermore, in this study, we identified 254 metabolic genes that are differentially expressed in CRC cells co-cultured with *Fn*. This indicates the profound impact microbes have on CRC metabolism and treatment responses.

Transcriptomics of colorectal HCT116 cells treated with *Fn* (GSE141805) was used with flux balance analysis (FBA) to generate a new condition-specific metabolic vector that represented the metabolic effects *Fn* had on HCT116 cells.^46^ This was used as described above to generate joint profiles with the 110 individual drug treatment metabolic profiles and predict drug-bacteria synergy scores. We repeated this process with transcriptomic data of HT29 cells treated with *Fn* (GSE90944).^47^ For two-drug combinations, the presence of *Fn* significantly altered predicted responses relative to predictions without *Fn* (p ≪ 0.05).

Most drugs were not predicted to be synergistic with *Fn* (Fig. 3c). The predictions were compared to a study where CRC patients with *Fn* presence were treated with capecitabine and a study on patient-derived *Fn* samples combined with fluorouracil.^48,49^ Patients treated with capecitabine showed high relapse rates, which correlates with our prediction of capecitabine not being a synergistic combination with *Fn*. Alternatively, fluorouracil was predicted to be strongly synergistic, consistent with its effectiveness in treating colon cancer.^49^

**Fig. 3.**
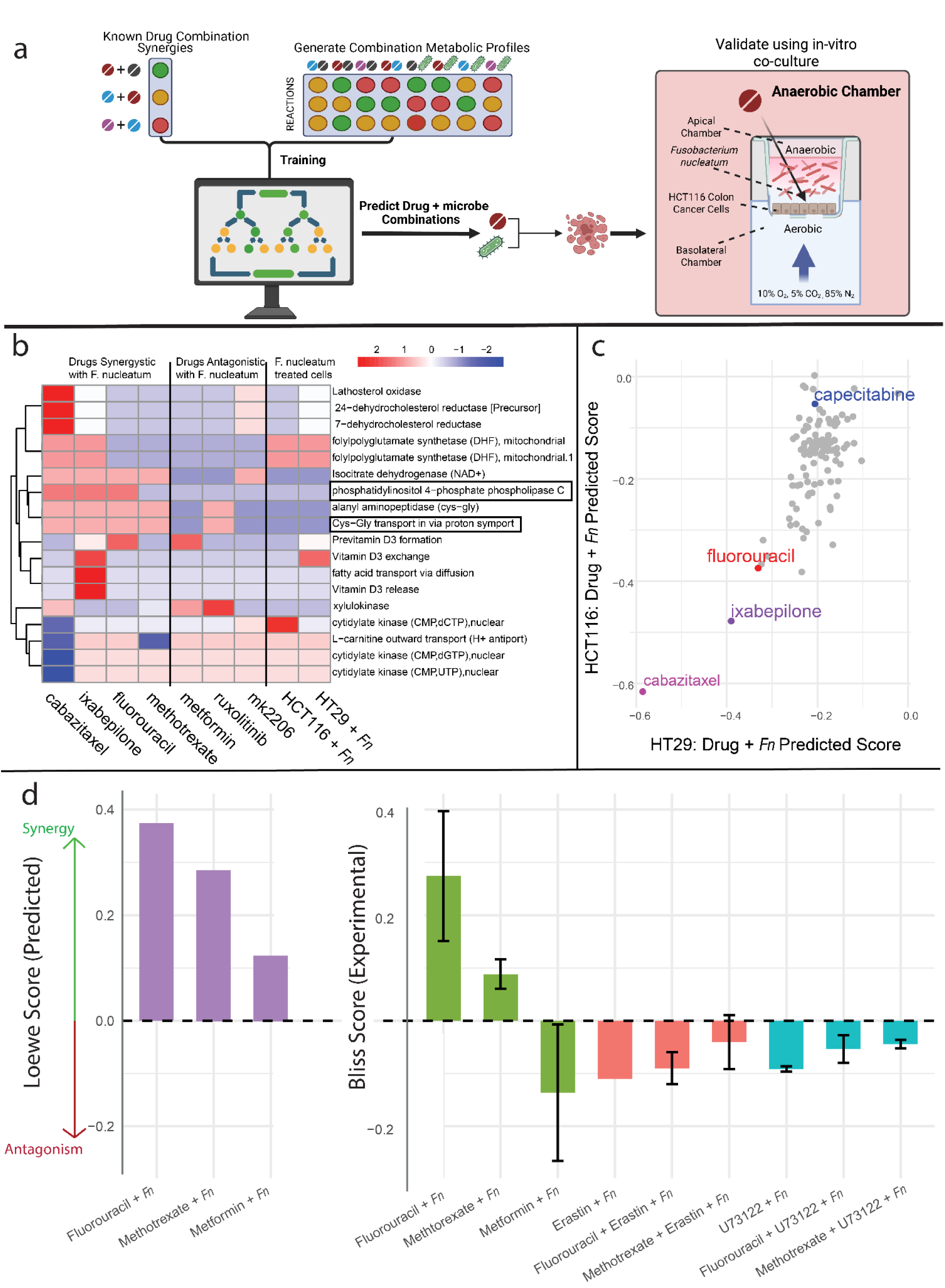
Modeling and experimental validation of *Fn*–drug interactions. a. Experimental setup for HCT116 and Fn co-culture. **(a)** An oxygen-containing carrier gas is introduced into the lower (basolateral) chamber of the Transwell system. As the gas diffuses upward, oxygen is progressively consumed by the HCT116 cells, resulting in an oxygen-depleted carrier gas reaching the apical compartment. This oxygen gradient establishes a hypoxic to anaerobic microenvironment within the upper Transwell, thereby enabling anaerobic bacteria to grow normally and exhibit infection behaviors that closely resemble those observed under physiological conditions. This setup enables a viable co-culture of both cell types by simultaneously fulfilling their oxygen needs*. Figure created using BioRender.com.* **(b)** Flux through top features for drugs synergistic and antagonistic with Fn. Shown drugs were selected based on being validated experimentally (fluorouracil, methotrexate, and metformin) or strong synergistic predictions (cabazitaxel and ixabepilone), or strong antagonistic predictions (axitinib and aminolevulinic acid). Columns for the flux of each cell line treated with only Fn are shown, as well. Feature importance was measured by how much each predictor reduces mean squared error (MSE) when used for splitting, averaged across all trees in the ensemble. Features that contribute more to reducing error were ranked as more important. The top 25 features are shown here. Boxed reactions were targeted experimentally. **(c)** Comparison of predicted Loewe scores for single drugs combined with Fn. Outputs vary by cell line (HT29 or HCT116). More negative values indicate synergy **(d)** Predicted synergy scores (Loewe) were multiplied by –1 to align their directionality with experimental measurements, but they still represent Loewe synergy values. For experimental assays, HCT116 cells were co-cultured with Fn for 24 hours prior to drug exposure. Cells were then treated for an additional 24 hours with 40 µM fluorouracil, metformin, or methotrexate; 10 µM erastin; or 1 µM U73122. Cell viability was measured using Cell Titer-Glo, and viability values were converted to Bliss synergy scores.

The drug predicted to be most synergistic in both cell lines was cabazitaxel, similar to predictions made for two drug combinations without *Fn*. The second most synergistic drug for both cell lines was predicted to be ixabepilone. Ixabepilone is a taxane commonly used to treat breast cancer in patients who don’t respond to docetaxel or paclitaxel.^50^ It is also known to demonstrate synergy with various other drugs, including capecitabine and brivanib.^51^ Interestingly, the model predicted ixabepilone to be involved in many of the most synergistic combinations both with and without *Fn* (Fig. 3c).

Three of the predicted combinations were validated *in vitro* (Fig. 3d) using a co-culture system that involves an asymmetric gas setup with both aerobic and anaerobic culture conditions, to allow both the HCT116 cell monolayer and *Fn* to grow effectively (Fig. 3a).^52^ This low number of samples was due to the complexity of the co-culture experimental system, which requires a low throughput and careful optimization. Despite these restraints, this modeling system is a much more accurate recapitalization of the tumor microenvironment compared to higher throughput methods. This allows us to assess the predictive ability and physiological relevance of the model’s predictions. Top candidates to test were prioritized from initial experiments without *Fn* (Fig. 2a). Fluorouracil was tested, given its clinical relevance for CRC, and as it was predicted to be synergistic in the presence of *Fn*.^49^ Metformin was selected to represent the results of a neutral (not synergistic) combination. Methotrexate was tested for its novelty in this setting, which has been applied to colon cancer, but it is not often a first-line treatment.^53^ Additionally, methotrexate is known to alter the microbiome in patients, but has not been studied in relation to *Fn.*^54^ Initially, cabazitaxel was applied as well. However, due to poor solubility, cabazitaxel was incompatible with the co-culture system.

The model was optimized to predict Loewe scores, whereas the *Fn* experiments resulted in Bliss scores. Therefore, the raw scores from the model’s prediction and the experiments cannot be directly compared, as the model predicts Loewe scores and the experiment results are in Bliss scores. We used Bliss independence for the experimental analysis because the drugs were screened at fixed concentrations with the bacteria determined by their individual dose–response profiles, making the Bliss framework more appropriate for this design. In contrast, the model uses Loewe additivity, which defines synergy relative to dose additivity of the same compound and is more suitable for evaluating interactions computationally across continuous concentration spaces. Both fluorouracil and methotrexate demonstrated synergistic effects with *Fn in vitro*, which aligns with the predicted scores being highly synergistic (Fig. 3d). In contrast, metformin displayed antagonistic effects in the co-culture system, reducing efficacy in the presence of *Fn* (Fig. 3d). This is expected as the predicted score for metformin + *Fn* (-0.12) was ranked the 77th most effective out of 111 drugs, whereas fluorouracil + *Fn* ranked 4th and methotrexate + *Fn* ranked 14th. Therefore, the qualitative direction of synergy and relative effects of drugs in combination with *Fn* align with predicted efficacy.

### Metabolic Insights on *Fn* Treatments

The top features from the model were next used to find reactions that are predictive of drug-microbe synergy. Fig. 3b shows the top 25 features and their relative flux for treatments that are synergistic and antagonistic with *Fn*. The drugs demonstrate great variability from each other, and from cells treated with just *Fn,* between these top features.

One pattern that warranted further investigation was the increased flux of cysteine-glycine import and alanyl aminopeptidase (cysteine-glycine) activity in cells treated with fluorouracil and methotrexate, in contrast to the decreased activity of the same reactions in cells treated with *Fn* (Fig. 3b). To test if this reaction was related to fluorouracil and methotrexate anti-tumor activity, the cells in co-culture were treated with these individual drugs and erastin. Erastin is a small molecule that inhibits cysteine import via SLC7A11 and accelerates oxidative stress.^55^ Therefore, it was hypothesized that erastin would disrupt cysteine transport and attenuate the synergistic effects observed with fluorouracil and methotrexate. Consistent with this, the Bliss score for fluorouracil and *Fn* decreased by 92% and the score for methotrexate and *Fn* decreased by 85% with the addition of erastin (Fig. 3d). Interestingly, erastin and *Fn* displayed weak synergistic effects (Bliss score = 0.13), suggesting that *Fn* may partially sensitize cells to oxidative stress when cysteine import is inhibited.

Additionally, three of the synergistic drugs have increased activity of phosphatidylinositol reactions, while the antagonistic drugs and the *Fn* treatments have decreased flux through this reaction (Fig. 3b). Inositol metabolism has a strong link to CRC, as inositol hexaphosphate has shown anticancer activity and enhances chemotherapy efficacy.^56,57^ This observation was further examined in vitro using the phospholipase C inhibitor U73122. Treatment of HCT116 cells with U73122 in the presence of *Fn* resulted in minimal synergy (Fig. 3d). However, the addition of fluorouracil or methotrexate to the U73122–*Fn* treatment led to weak antagonistic interactions (Fig. 3d).

These results suggest that the synergistic effect of fluorouracil and methotrexate with *Fn* is partly mediated through perturbations in cysteine metabolism and phosphatidylinositol metabolism, and concurrent inhibition of these pathways reduces their synergistic effect based on the Bliss independence model of drug interactions.

### Predictions of Drug Combinations and Immunotherapy with Other Microbes

The connection between microbes and cancer therapeutics is being increasingly investigated, particularly in CRC.^11,20^ However, the gut microbial community is incredibly diverse between patients, which makes it difficult to pinpoint the exact impact of individual microbes.^11^

Therefore, in addition to *Fn*, four other microbial species were added to the model for drug combination predictions. These microbes were *Aeromonas veronii* (*A. veronii*, *Av*)*, Escherichia coli* (*E. coli, Ec*)*, Enterococcus gallinarum* (*E. gallinarum*, *Eg*), and *Streptococcus gallolyticus* (*S. gallolyticus*, *Sg*).^58–60^ *Sg* and *Ec* were selected for their strong ties to CRC. ^11^ Additionally, we included *Eg* (a potential biotherapeutic) and *Av* (a gut pathogen) to analyze the diversity of microbial influences.^59,61^

These microbes were also selected because they had a transcriptional impact on the HT29 cells in co-culture that could be mapped to the RECON1 metabolic model. However, the effects of microbial co-culture on HT29 cells were considerably weaker than those of drug treatments, necessitating a lower z-score threshold for identifying differentially expressed genes (±1 for drug treatments vs. ±0.2 for bacterial co-culture). Since the residence time of these microbes in the colon are much longer than drugs, we hypothesized that these small transcriptional changes might still have a significant metabolic impact.

Microbe-conditioned predictions with two drug combinations revealed a consistent, significant reduction in drug synergy across all tested microbes, with *Fn* exerting the strongest effect, followed by *Eg*, *Sg*, *Av*, and *Ec* (FDR < 0.001 for all). Additionally, predicted scores for each microbe–drug combination differed significantly across all pairings, except for *Eg* vs. *Av* and *Sg* vs. *Av* (Fig. 4a). Additionally, Cabazitaxel, ixabepilone, and fluorouracil consistently ranked among the top five most synergistic combinations across all bacterial treatments. Notably, drug interaction outcomes varied both across microbial strains and between the two colorectal cancer cell lines (Fig. 4b, Supp. Fig. 5b. These differences likely reflect cell line–specific biological characteristics, such as distinct genetic backgrounds or metabolic states, and may indicate differential metabolic modulation by individual microbial species.^62^

**Fig. 4.**
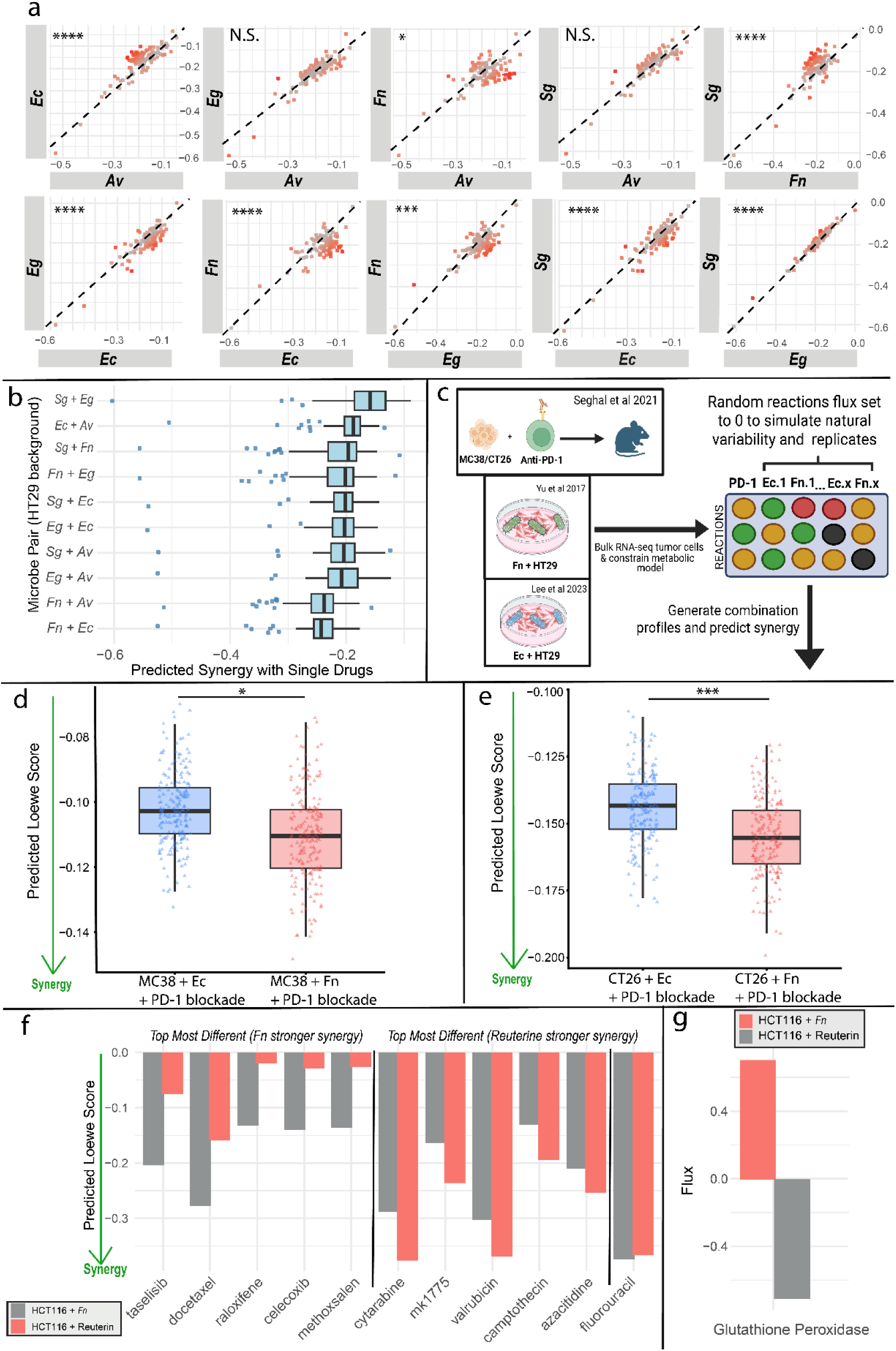
Predicted effects of individual and combined microbial treatments on drug synergy and PD-1 blockade. **(a)** Scatter plot matrix of predicted drug synergy scores of the drug combined with the given microorganism. P-value calculated using paired t-test with Bonferroni correction. The dotted line represents perfect correlation; therefore, values above the dotted line are drugs predicted to have a higher Loewe score in the y-axis microbe, and values below the dotted line are drugs predicted to have a higher score in the x-axis microbe. Red points represent the drugs that have predicted scores that are the most different under the different bacterial treatments. P-values (left to right). Top row: <0.001, >0.999, 0.021, 0.07, <0.001; bottom row: <0.001, <0.001, 0.001, <0.001, <0.001. **(b)** Predicted synergy of individual drugs with two microbe combinations. More negative values indicate synergy. The drug with the most negative predicted score for all microbe treatments is cabazitaxel. **(c)** Schematic methods used to produce data for Fig 4d and Fig 4e. Transcriptomics from anti-PD-1 treated MC38 and CT26 colon cancer cells injected into mice and HT29 cells treated with *Fn* or *Ec* were used to constrain a metabolic model. Then, simulated combined flux profiles were used to predict synergy of microbes and anti-PD-1 therapy. *Figure created using BioRender.com.* (**d-e)** Differences in predicted synergy of *Ec* and anti-PD-1 therapy or *Fn* and anti-PD-1 therapy on MC38 or CT26 colon cancer cells. Welch’s two-sample *t*-test was used to compare 200 simulated replicates per condition for each cell line (CT26, *p* = 4.39 × 10⁻¹⁴; MC38, *p* = 3.08 × 10⁻¹⁰). **(f)** Predicted Loewe scores of single drugs in combination with either *Fn* or reuterin on HCT116 cells. The plot highlights the top five drugs for which *Fn* produced stronger predicted synergy and the top five for which reuterin produced stronger predicted synergy. Fluorouracil is bolded because it was validated experimentally. More synergistic scores are more negative. **(g)** Z-scored metabolic flux through glutathione peroxidase reaction in HCT116 cells treated with either *Fn* or reuterin. More positive values indicate higher flux relative to the reaction mean.

Since the colon microbiome is rarely a monoculture, we next simulated the effects of pairwise combinations of microbes with and without drug treatments (Fig. 4b, Supp. Fig. 5a-b). Some combinations of microbes showed strong synergistic effects on drugs compared to individual microbes alone - especially combinations involving *Av* and *Fn*, and *Ec* and *Fn*. As individual ‘treatments’, each microbe was also weakly synergistic with the others, suggesting mild inhibitory activity against the CRC cell lines, although there were strong differences between the two cell lines for the combination of *Ec* and *Fn*.

Comparing flux through top features for microbe-treated HT29 cells, it is apparent that *Fn* and *Ec* have distinct profiles (Supp. Fig. 5a). In particular, *Fn-treated* HT29 cells have a strong increase in lathosterol oxidase, dehydrocholesterol reductase, and vitamin D metabolism compared to the other microbial treatments. *Ec*, on the other hand, induced increases in phospholipase C, alanyl aminopeptidase, cysteine-glycine transport, and decreased glycine reversible transport and L-carnitine outward transport. Interestingly, flux through top features for HT29 cells treated with *Fn* was relatively different from the flux of HCT116 cells treated with *Fn*. In particular, *Fn* seemed to have a less profound impact on HCT116 cells, inducing only 106 total DEGs in HCT116 cells compared to 245 in HT29 cells. Importantly, the transcriptomics data used were sourced from unique studies, and different conditions could explain this discrepancy, as predictions from the two cell lines treated with *Fn* correlated strongly (Fig. 3c).

Although the model was trained exclusively on chemotherapy drug combinations, we next evaluated its applicability to more contemporary colorectal cancer (CRC) treatment strategies, including immunotherapy. Transcriptomic data from Sehgal *et al.* were used to generate metabolic flux profiles corresponding to αPD-1 treatment conditions.^63^ To benchmark model performance in this context, we predicted the combination synergy between αPD-1 therapy and microbial coculture. *Fn* has been previously associated with improved responses to anti–PD-1 therapy compared with *Ec* K-12–derived strains.^64,65^ Consistent with these experimental observations, the model predicted significantly greater synergy between *Fn* and αPD-1 therapy compared with *Ec* and αPD-1 therapy across two CRC cell line models (CT26, *p* = 4.39 × 10⁻¹⁴; MC38, *p* = 3.08 × 10⁻¹⁰; Fig. 4c-e). Collectively, these results demonstrate that the model generalizes beyond conventional chemotherapy combinations and can capture metabolic interactions in more complex immunotherapy-associated microenvironments.

Finally, we simulated the effects of probiotic treatment. Ruterin, a metabolite produced by the probiotic strain *Limosilactobacillus reuteri* (*Lr*), suppresses tumor growth by disrupting redox balance in colon cancer cells. Additionally, *Lr* demonstrates synergism with fluorouracil.^66^ In line with these experimental observations, we predicted *Lr* and fluorouracil to be strongly synergistic (Fig. 4f). Fluorouracil was the 7th most synergistic drug in combination with *Lr* of the 110 drugs tested. Across all 110 drugs, the predicted synergy profiles for *Fn* versus reuterine were significantly different (*p* = 4.85 × 10⁻¹²). Furthermore, the effect of *Lr* on colon cancer has been linked to an increase in oxidative stress by depleting glutathione.^66^ Likewise, the flux through glutathione peroxidase was decreased in reuterine-treated cells compared to *Fn*-treated cells (Fig. 4g). This demonstrates the model’s accuracy and generalizability across bacterial species.

## Discussion

Systems biology approaches are necessary to comprehend the metabolic variation and crosstalk within tumor microenvironments (TME). The combinatorial complexity involving the interaction between specific microbial strains, drug regimens, and tumor metabolism makes it challenging to model the TME *in vitro* and renders exhaustive experimental screening impossible. We present a generalizable machine-learning framework, OMG-ML, that models interactions across diverse microbial combinations, novel drugs and drug combinations, as well as probiotic interventions, enabling the systematic identification of high-impact therapeutic strategies. This represents a unique framework to tailor drug regimens for cancer therapy based on microbial species.

The OMG-ML pipeline demonstrates robust predictive performance for chemotherapy combinations, capturing both synergistic and antagonistic interactions across colorectal cancer cell lines. Model performance was verified using cross-validation, independent test sets, experimental results, and comparisons to similar and clinical studies (Fig. 2a-e). Model predictions were further supported by *in vitro* experiments, confirming the predicted efficacy of specific drug combinations such as cabazitaxel and megestrol. The combination of cabazitaxel and megestrol underscores the power of machine learning in predicting drug synergies, as neither taxanes nor progesterone are commonly used alone or together for colon cancer treatment.

A key advantage of combining machine learning with FBA is the ability to extract the most influential features driving the model’s predictions. These features can offer mechanistic insights into why certain drug combinations are predicted to be synergistic. These reactions may represent critical metabolic vulnerabilities in CRC and, therefore, serve as potential therapeutic targets. We identified increased phosphoinositol metabolism and cysteine transport in drugs that were effective in the presence of *Fn* and inhibited these pathways to identify if these effects existed *in vitro*. Lathosterol oxidase and many other enzymes in the top features have strong links to redox signaling and ferroptosis. These include 7- and 24-dehydrocholesterol reductase, folylpolyglutamate synthase, and cysteine-glycine transport (Fig. 3a-b).^67–69^ These pathways are increasingly recognized as promising therapeutic targets in CRC.^70^

Beyond *Fn*, the five additional bacterial species and the reuterin treatment revealed several notable patterns. Reuterin’s predicted effects were consistent with prior experimental findings, including reduced glutathione peroxidase activity and enhanced synergy with fluorouracil.^66^ Across microbe–drug combinations, 8 of the 10 pairwise comparisons showed significantly different predicted responses, underscoring the extent to which microbial context can reshape drug efficacy. Moreover, introducing multiple microbes simultaneously (microbe–microbe–drug predictions) consistently shifted responses toward greater synergy, suggesting that microbial diversity within the tumor microenvironment may potentiate therapeutic effectiveness. Additionally, in all five tested bacterial species, cabazitaxel, ixabepilone, and fluorouracil ranked in the top five most effective drugs, highlighting their general strength in unique microbiomes.

Our model can also predict beyond traditional chemotherapies, as illustrated in Fig 4c-e where the model accurately predicted PD-1 blockade to be more synergistic with *Fn* than *Ec*. This result may seem counterintuitive, but several in vivo and patient studies have found Fn levels to be associated with improve immune therapy responses (PD-1 and PDL-1 blockade).^64,65,71^ However, the relationship between *Fn* and immunotherapy is not well understood and some studies contradict *Fn* improving treatment outcomes.^72,73^ As more research emerges on the dynamics between microbes and immunotherapy, our model’s ability to recapitulate this relationship and predict the outcomes can be further validated and fine-tuned.

Our model stands out from other similar models by its scalability, accuracy, interpretability of pathways of interest, and its connection to the tumor microenvironment. Together, these results illustrate that integrating metabolic modeling, machine learning, and microbial context can reveal novel therapeutic opportunities and mechanistic insights for colorectal cancer. Future studies incorporating larger patient datasets and metabolomic validation will be necessary to confirm these trends and refine the predictive capacity of flux-based models in clinical settings.

## Methods

### Metabolic Modeling

Transcriptomic profiles of HT29 and HCT116 colorectal cancer (CRC) cells treated with a total of 110 drugs were obtained from the LINCS data portal, comprising 95 treatments in HT29 cells and 15 in HCT116 cells.^38^ Due to limited treatment data for HCT116, an additional condition was generated using differentially expressed genes (DEGs) from untreated HCT116 cells compared to HT29 cells, sourced from the Cancer Cell Line Encyclopedia (CCLE). DEGs from co-cultures of HCT116 and HT29 cells with *Fusobacterium nucleatum* were obtained from the Gene Expression Omnibus (GEO; GSE141805; GSE90944).

Each treatment condition’s DEGs were integrated into the RECON1 or RECON3D genome-scale metabolic model (GEM) to generate condition-specific metabolic models. From drug treatments, genes with z-scores > 1 or < –1 were considered significantly up- or downregulated, respectively. Whereas the threshold was set to z-scores > 0.2 or < –0.2 for microbial co-cultures. Integration into RECON1 was performed using a linear optimization-based implementation of the integrative metabolic analysis tool (iMAT), which maps transcriptomic changes to shifts in metabolic flux.^74,75^ The iMAT algorithm was parameterized using kappa = 10⁻², rho = 10², and epsilon = 1, where kappa and rho weight the penalties for “off” and “on” reactions, and epsilon specifies the minimum allowable flux through “on” reactions. All reported analyses employed the biomass objective function. Although other objective functions, such as maximizing ATP production, were evaluated, none provided better performance in 10-fold cross validation than the biomass objective.

This process generated 113 individual metabolic flux profiles. Joint flux profiles were generated for each drug combination using four key features: sigma score, delta scores, and entropy. These scores capture shared (sigma) and unique (delta) metabolic changes between drugs. Sigma and delta scores were derived from binarized flux profiles that indicated whether each metabolic reaction was more or less active compared to baseline. For each condition (drug or co-culture), the reactions were marked as +1, -1, or 0 depending on the direction of flux change. These binarized profiles were then compared across drug pairs. Sigma scores reflected reactions affected by both drugs (shared effects), while delta scores captured reactions uniquely influenced by one drug but not the other (distinct effects). Cumulative entropy was calculated using un-binarized flux profiles. First, we computed the metabolic entropy for each condition, which reflects the variability of fluxes across all reactions. Higher entropy indicates a more widespread or less targeted metabolic effect. Then, for each drug combination, we summarized these entropy values by taking both their mean and total sum – yielding two features that describe the overall extent of metabolic disruption caused by the combination.

To model synergism with anti-PD-1 therapy, transcriptomics from MC38 and CT26 cells injected into mice along with anti-PD-1 blockade were sourced from Seghal et al.^63^ Differential gene expression data were originally generated by comparing treated and untreated tumor samples. To integrate this data into our modeling pipeline, differentially expressed genes (DEGs) between these tumors and HT29 cells were identified and used to constrain metabolic models and calculate reaction fluxes. To account for biological variability, metabolic flux profiles from *Ec* and *Fn* coculture simulations were subjected to reaction dropout analysis. Specifically, for each treatment condition, 20% of reaction fluxes were randomly set to zero, and this procedure was repeated 200 times to generate replicate flux distributions. These simulated replicates were then used to perform statistical comparisons (two-sided Welch’s *t*-tests) between the two anti–PD-1 plus microbial coculture conditions to assess metabolic synergy.

### Comparing Metabolic Models

The model was trained using the RECON1 metabolic network, which showed higher accuracy on validation data than RECON3D. On independent test data from synergyxDB, predictions from the RECON1-trained model correlated significantly with observed responses in both HT29 and HCT116 cells (Pearson’s r = 0.79 and 0.51, respectively; Fig. 2d–e). The RECON3D-trained model also correlated significantly with the same test data (HT29: r = 0.62; HCT116: r = 0.52). AUC values followed a similar pattern: for HT29 and HCT116 cells, the RECON1-trained model achieved 0.77 and 0.91, respectively, compared to 0.75 and 0.83 for the RECON3D-trained model. On new in vitro predictions, the RECON1-trained model performed slightly better (r = 0.48) than the RECON3D-trained model (r = 0.47). However, the RECON3D model performed slightly better in 10-fold cross-validation (r=0.57, p-val < 2.2e-308; synergy AUC = 0.81; antagonism AUC = 0.75). Despite these slight performance differences, predictions from the two models correlated closely for both cell lines (HT29: r = 0.81; HCT116: r = 0.79; Fig. 7a-b). Interestingly, although the top features identified by each model were entirely distinct (Fig. 7c), the distribution of metabolic subsystems among these features was similar (Fig. 7d), particularly for extracellular transport reactions and inositol metabolism, highlighting the importance of these pathways.

### Machine Learning Prediction

In this study, we developed OMG-ML to predict synergistic colon cancer drug combinations, incorporating metabolic influences from the tumor microenvironment. OMG-ML was built in MATLAB using the fitrensemble bagging algorithm. This algorithm constructs multiple decision trees, each trained on a different bootstrap sample of the original data. The final prediction is obtained by averaging the outputs of all learners. In our implementation, fitrensemble was used with default parameters, which include decision trees as base learners with a maximum number of 10 splits per tree (i.e., MaxNumSplits = 10), a maximum of 100 learners (NumLearningCycles = 100), and no pruning. The input to the model consisted of binarized flux profiles representing individual condition-specific metabolic activity.

6514 known Loewe synergy scores were used in training (HT29 N = 3256, HCT116 N = 3258). Because only 15 of the drug treatments in the LINCS transcriptomic dataset overlapped with the 6514 drug combinations used to train the model for HCT116 cells, we incorporated an additional baseline condition to better represent HCT116-specific metabolism. This condition was generated by identifying differentially expressed genes (DEGs) between untreated HCT116 and HT29 cells, using RNA-seq data from the Cancer Cell Line Encyclopedia (CCLE). This allowed for training on all available Loewe scores in SynergyxDB and for predictions to be made based on cell type.

To benchmark the model’s performance, we also trained models using transcriptomic input, including a bagged ensemble of decision trees (random-forest–like) and a gradient-boosted regression model (GBM). These models use drug-induced gene-expression signatures to predict combination responses and provide a direct comparison to the flux-based framework (Supplementary Fig. 4a. On 10-fold cross validation these models resulted in an AUC of 0.80 and 0.81, respectively, which is similar to the flux-based model performance. However, when testing on a withheld dataset, they performed worse with Pearson’s r of 0.55 and 0.52 for HT29 data and 0.40 and 0.39 for HCT116 data (Supplementary Fig. 4.b-e). Comparatively, the flux-based model achieved Pearson’s r of 0.79 and 0.51 on the same test set (Fig. 2c-d). Furthermore, the flux-based model’s performance was assessed using a permutation test, in which the data labels were randomly shuffled 1000 times and the resulting null distribution of R^2 values was compared against the model’s true R^2 to evaluate statistical significance. The p-value from this test was very significant (p-value = 0.000999), indicating the model learned a meaningful biological or statistical signal.

We also manually assessed if the fluxes match known effects of drug treatment. For example, allopurinol treatment led to decreased flux through its target xanthine oxidase.^76^ However, not all drugs were as clean-cut. Specifically, floxuridine and capecitabine had increased flux through thymidylate synthase despite being inhibitors of this enzyme.^77,78^ Despite this, the other drugs that target these enzymes, fluorouracil, methotrexate, and pemetrexed induced an expected decrease in flux.^79–81^ Likewise, aminolevulinic acid caused a decrease in flux through aminolevulate synthase, although this direct inhibitory link has not yet been observed in colon cancer.^82^ Interestingly, mercaptopurine and thioguanine treatment caused a decrease in hypoxanthine phosphoribosyltransferase, which is an essential enzyme in the bioactivation of these drugs.^83,84^

Feature importance was assessed by evaluating the reduction in mean squared error (MSE) attributed to each predictor across the entire ensemble model. Specifically, for each decision tree within the ensemble, the algorithm tracked how much each feature contributed to decreasing MSE at each split point. These individual reductions were then averaged across all trees to obtain an overall importance score for each feature. Features that consistently led to greater error reduction were assigned higher importance, reflecting their stronger influence on the model’s predictive accuracy.

### Transcriptomic-Based Prediction

The flux-based model was compared to a model developed using transcriptomic data as input. Differentially expressed genes (DEGs) were the same used to determine flux (thresholds of |log2FC| ≥ 1]). The DEGs were converted into features using a similar style to OMG-ML by generating three feature classes. The first class modeled the combined transcriptional response under independent action by averaging the two drug signatures using the equation below (DEG_A_ and DEG_B_ are the expression of two specific genes in the drug signature):

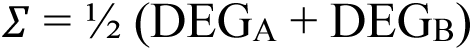

The second feature captured the disagreement between the two perturbations:

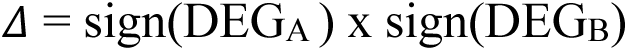

The last feature was entropy, which characterized the breadth of transcriptional disruption. An entropy metric for each drug was computed, and gene-level magnitudes were normalized to form a probability distribution, pi, and entropy was calculated as:

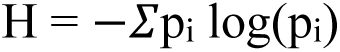

For each drug pair, the *Σ*, *Δ* , and H features were concatenated to form a joint feature vector that represents the predicted combined transcriptional state of the two drugs. These feature vectors were used to train both a random forest model and a gradient boosting model. Synergy labels were obtained from SynergyXDB, and model performance was evaluated using 10-fold cross-validation and a withheld independent test set.

### Experimental Validation

#### Two Drug Combinations

HCT116 cells were cultured in Dulbecco’s Modified Eagle’s Medium and were treated individually with metformin, cabazitaxel, megestrol acetate, sorafenib, erlotinib, gemcitabine, and methotrexate. They were also treated with combinations of cabazitaxel and megestrol with each of the aforementioned drugs (11 total combinations). The cells were treated at doses of 0.9, 5, 10, 20, and 40 μM. Cell viability was assessed using Cell Titer Glo staining, and results were used as input for SynergyFinder to derive Loewe Scores.

#### Single Drug and *Fn*

In the colon, intestinal cells are exposed to oxygen from the blood supply, but microorganisms are not. This creates an oxygen gradient that cannot be recapitulated by simple co-culture systems. Therefore, we used a system where oxygen is pumped to CRC cells plated on a basal membrane of a 24-well plate. In the upper chamber of the well, *Fn* cells are seeded. The rate of oxygen flow to the CRC cells is roughly equivalent to oxygen uptake, making the upper chamber anaerobic. This provides an optimal environment for both cell types to survive long term, whereas in a simple co-culture setup, *Fn* would die after several hours, causing additional stress to the CRC cells. While this setup is one of the most accurate ways to recapitulate the CRC TME, it is limited by the size of the plate that is compatible (maximum of 24 wells). This restricted our possible predicted drug treatments to validate. We tested 4 total drugs in combination with *Fn*: fluorouracil, methotrexate, metformin, and cabazitaxel. However, cabazitaxel testing was not completed due to solubility restraints. HCT116 cells were co-cultured with *Fn* for 24 hours prior to treatment with 40 uM of each drug and then incubated with both *Fn* and the drug for an additional 24 hours. Cell Titer Glo was used to assess viability and determine synergy scores. HCT116 cells were co-treated with fluorouracil or methotrexate and the pathway inhibitors erastin (10 µM) or U73122 (1 µM) in the presence of Fn. All combinations were applied 24 hours prior to viability measurement.

#### Bacterial culture conditions

*Fusobacterium nucleatum* (ATCC 23726, https://www.atcc.org/) was initially inoculated from 80 °C glycerol stock into Columbia broth (CB) medium (BD, 294420) 48 h before experimentation and cultured under anaerobic conditions (10% CO₂, 5% H₂, and 85% N₂, Welding Supply Corp.) at 37 °C using a vinyl anaerobic chamber (COY Type A).^85,86,87^ Once the culture reached turbidity (OD_600_=0.5), a secondary inoculation was performed into the co-culture medium (CCM) one day before the start of the experiment, with a dilution fold of 1:50.^52^ The bacteria were grown under the same anaerobic conditions until reaching the mid-logarithmic growth phase (OD_600_=0.5), when cell viability was optimal for subsequent assays.

#### Collagen IV coating of Transwell Inserts

Collagen IV stock solution (1 mg/mL) was prepared by dissolving 5 mg of lyophilized collagen IV (Sigma, C5533-5mg) in 5 mL of cold 100 mM acetic acid (Fisher Scientific, A38-500).^88^ The collagen solution was rotated at 4 °C for a minimum of 5 h to allow complete solubilization, aliquoted under sterile conditions, and stored at –80 °C until use.

For Transwell coating, collagen IV aliquots were thawed at 4 °C for at least 5 h. A working solution of 33 µg/mL was prepared by diluting the stock solution in cold sterile water (Milli-Q, Sigma-Aldrich).^89^ Transwell plates (Avantor, 76313-906) were pre-cooled at –20 °C, and 100 µL of the collagen solution was added to each insert (10 µg/cm²).^90^ Plates were incubated on ice at 4°C for 30 min, followed by incubation at 37 °C for 4 h to allow collagen solidification. After coating, Transwells were used immediately or wrapped with parafilm and stored at 4 °C until use.

#### Preparation of the cell monolayer of HCT116 cells

HCT116 cells (ATCC CCL-247) were thawed from frozen stocks and expanded once at a 1:3 split ratio. When cultures reached ∼70% confluence, cells were detached with 0.25% trypsin–EDTA (Gibco, 25200056) for 2 min at 37 °C, neutralized with complete DMEM, and washed twice with sterile phosphate-buffered saline (PBS).^91^ Cells were resuspended at a final concentration of 2 × 10⁶ cells/mL. Transwell inserts (0.4 µm pore size, polycarbonate membrane; Corning) pre-coated with collagen IV were aspirated to remove residual coating solution, and 100 µL of the cell suspension was seeded into each apical chamber, resulting in 2 × 10⁵ cells per insert. Basolateral chambers were filled with 600 µL of DMEM supplemented with 10% heat-inactivated fetal bovine serum (FBS; Gibco, A5670701) and 1% penicillin–streptomycin (P/S, 10,000 U/mL, Gibco, 15140122). Co-cultures were maintained at 37 °C in a humidified incubator with 5% CO₂ for 4 days. The apical medium was refreshed every 48 h, while the basolateral medium was left undisturbed unless otherwise indicated.

#### Asymmetric anaerobic co-culture of HCT116 cells

The co-culture procedure was performed as previously described with minor modifications. Briefly, gas-permeable plates pre-filled with 600 µL of DMEM per well were secured onto the co-culture chamber using screws. The inlet and outlet ports were connected to a mixed gas stream (10% O₂, 5% CO₂, 85% N₂) at a final flow rate of 0.5 standard cubic feet per hour (SCFH), and magnetic stirring was applied to ensure uniform gas distribution. Transwell inserts containing confluent HCT116 monolayers were removed from the incubator, and the culture medium in the apical chamber was replaced with DMEM without antibiotics. Inserts were transferred into the anaerobic chamber and maintained under continuous gas flow for 3 h to allow equilibration of the atmosphere.

*Fn* previously passaged into CCM medium was harvested, adjusted to the desired concentration, and diluted to achieve a multiplicity of infection (MOI) of 0.1. The apical chamber medium was replaced with the diluted bacterial suspension, and co-cultures were incubated for 24 h under anaerobic conditions. At the 24-h time point, chemotherapeutic agents (40 mM final concentration) were added to each Transwell (inhibitors or stimulators added at the indicated concentrations), followed by an additional 24 h of incubation before downstream analyses. This setup was replicated for the U73122 inhibitor (1 uM final concentration) in combination with 40 mM methotrexate or fluorouracil and 0.1 MOI *Fn*.

#### Verification of the cell monolayer viability

For cell viability verification, the CellTiter-Glo® 2.0 assay (Promega, G9241) was used. After co-culture, the cell monolayer was removed, and both the apical and basolateral chamber solutions were replaced with DMSO containing penicillin/streptomycin (P/S).^92^ The monolayer was incubated for 2 hr under this treatment. Following treatment, the cells were washed with PBS, and 100 µL of CellTiter-Glo® 2.0 reagent was added to the apical chamber. After 10 min of incubation, bioluminescence images were acquired using the Azure 300 Imaging System with auto-exposure mode.^93^ The bioluminescence intensity of the Transwell inserts was subsequently quantified using a SpectraMax M5 plate reader (Molecular Devices).^94^

After imaging, the Transwell inserts were washed once with PBS. To evaluate the dead cell ratio, co-staining with SYTO9 Green (Thermo Fisher, S34854) and ethidium bromide (1% solution, Fisher Scientific, BP1302-10) were performed, each at a 1:10,000 dilution. The staining solution was added to both the apical and basolateral chambers, and the monolayer was incubated for 10 min. Fluorescence images were then acquired using a Nikon ECLIPSE Ts2-FL microscope.^95^

### Data & Code Availability

All code necessary to train and test the model as well as generate figures provided in this text can be found in the following github repository. https://github.com/anniejs02/OMG_ML

## ACKNOWLEDGEMENTS

This work was supported by faculty start-up funds and Rogel Cancer Center from the University of Michigan (UM), R35GM137795 from the National Institute of General Medical Sciences, UM Endowment for Basic Sciences Accelerator Award, UM Research Scouts Award, and Michigan Drug Discovery Screening Grant to S.C. , and National Science Foundation Graduate Research Fellowship (NSF GRFP) to AB We thank Rupa Bhowmick, Margaret Reuter and Katherine Lev for feedback and research assistance.

## AUTHOR CONTRIBUTIONS

Conceptualization: SC, AB; Methodology: AB, ZP, AR, BB, RN, CHC; Modeling: AB, CHC, BB; Experiments: AB, ZP, RN; Supervision: SC, JL; Writing—original draft: AB, SC; Writing—review & editing: AB, JL, SC

## Supplementary Materials

**Supp. Fig. 1.**
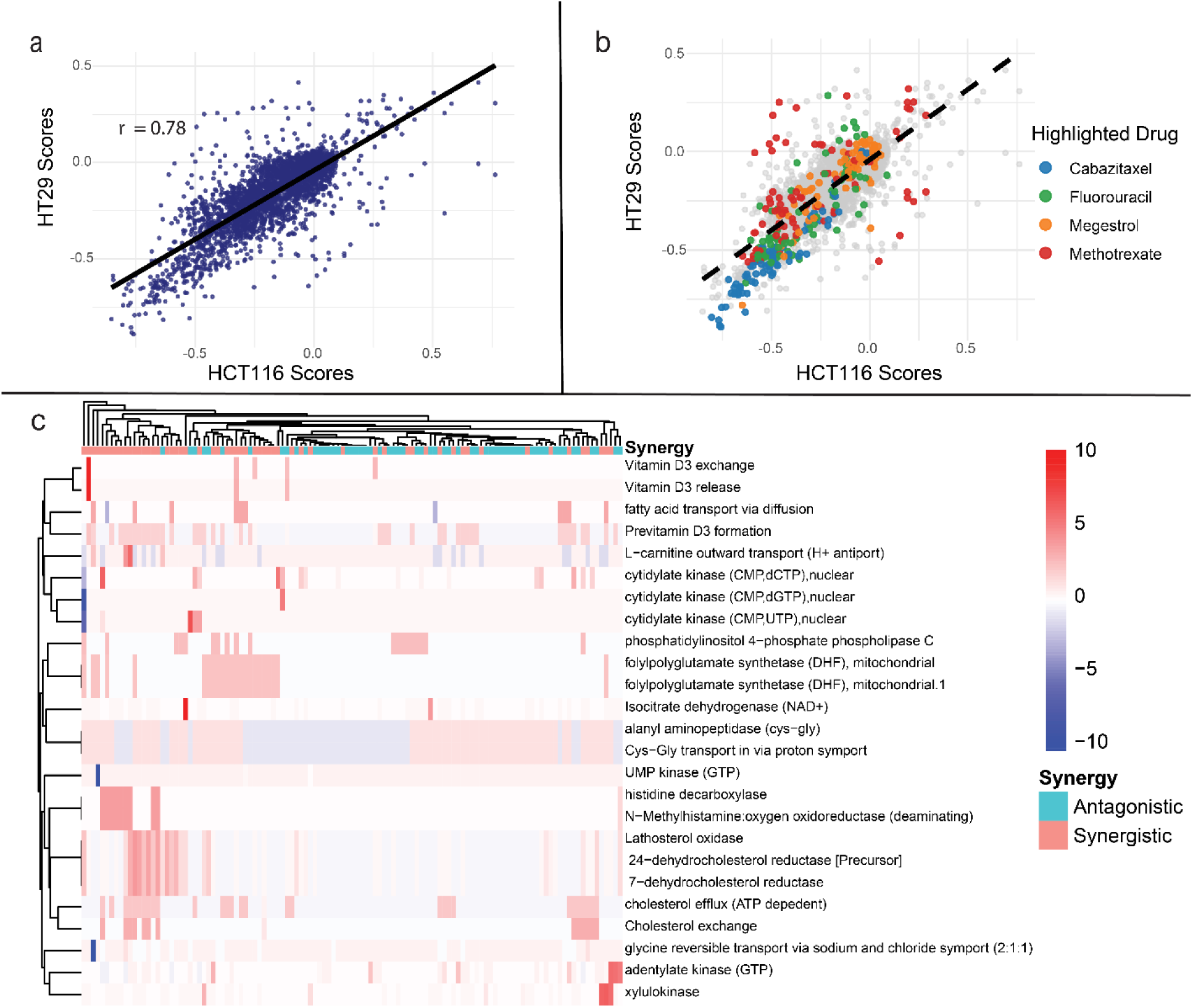
Training data distribution, patient predictions, and top feature importance. (a-b) *Correlation of known synergy scores from SynergyxDB for HT29 and HCT116 cells with drugs of interest highlighted*. **(c)** *Z-score normalized flux through top features for all drug treatments. Synergy and Antagonism labels were created by taking all combinations in which each drug was present and averaging the predicted scores. If the averaged score fell in the bottom 50%, the drug was considered synergistic, and if it fell in the top 50% it was labeled antagonistic*

**Supplementary Fig. 2.**
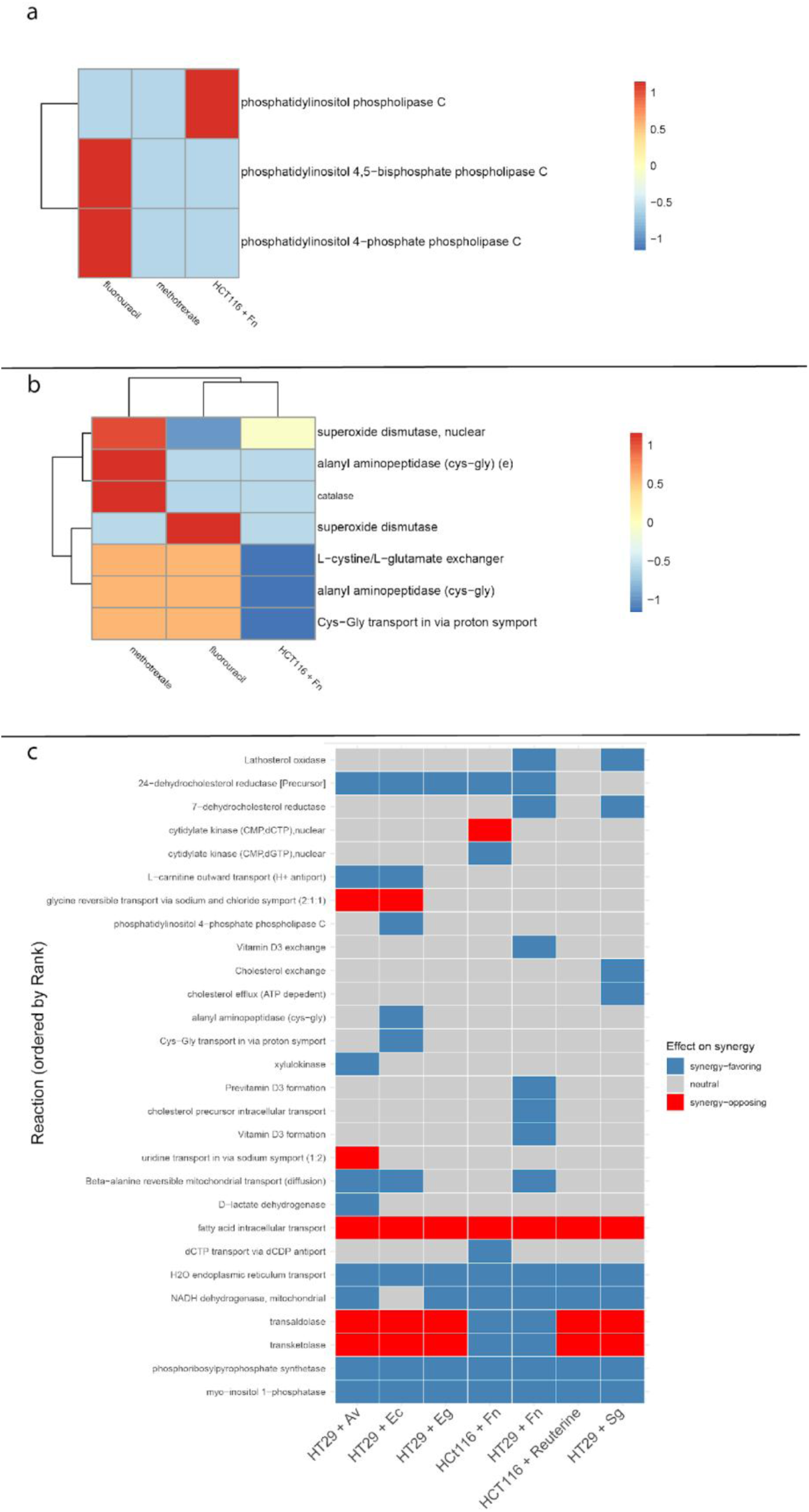
Flux through targeted pathways and microbial effects. **(a)** Flux through non-zero phospholipase C reactions for HCT116 cells treated with methotrexate, fluorouracil, or Fn. Values are z score normalized. **(b)** Flux through non-zero reactions related to ferroptosis and erastin’s effects for HCT116 cells treated with methotrexate, fluorouracil, or Fn. Values are z score normalized. **(c)** Alignment of microbe-induced fluxes with top synergy-associated reaction features. Heatmap showing whether each microbe’s predicted flux through the top model reactions pushes metabolic activity toward or away from the metabolic signatures associated with drug synergy. Each reaction (rows) is ordered by its feature importance rank, where Rank 1 represents the strongest predictor of interaction outcome. For each reaction, a Direction label was calculated (“pos” or “neg”), indicating whether increased flux through that reaction is associated with greater synergy (“pos”) or greater antagonism (“neg”) in drug–drug interaction scores. Microbe fluxes were multiplied by this Direction sign to produce a signed flux value, where positive values indicate flux that aligns with synergy-associated directions, negative values indicate flux that opposes them, and zero indicates no effect. Tiles are colored accordingly: blue (“synergy-favoring”), red (“synergy-opposing”), and grey (“neutral”). This visualization highlights which microbes recapitulate synergy-linked metabolic patterns and which shift cells into an antagonistic metabolic state.

**Supplementary Fig. 3.**
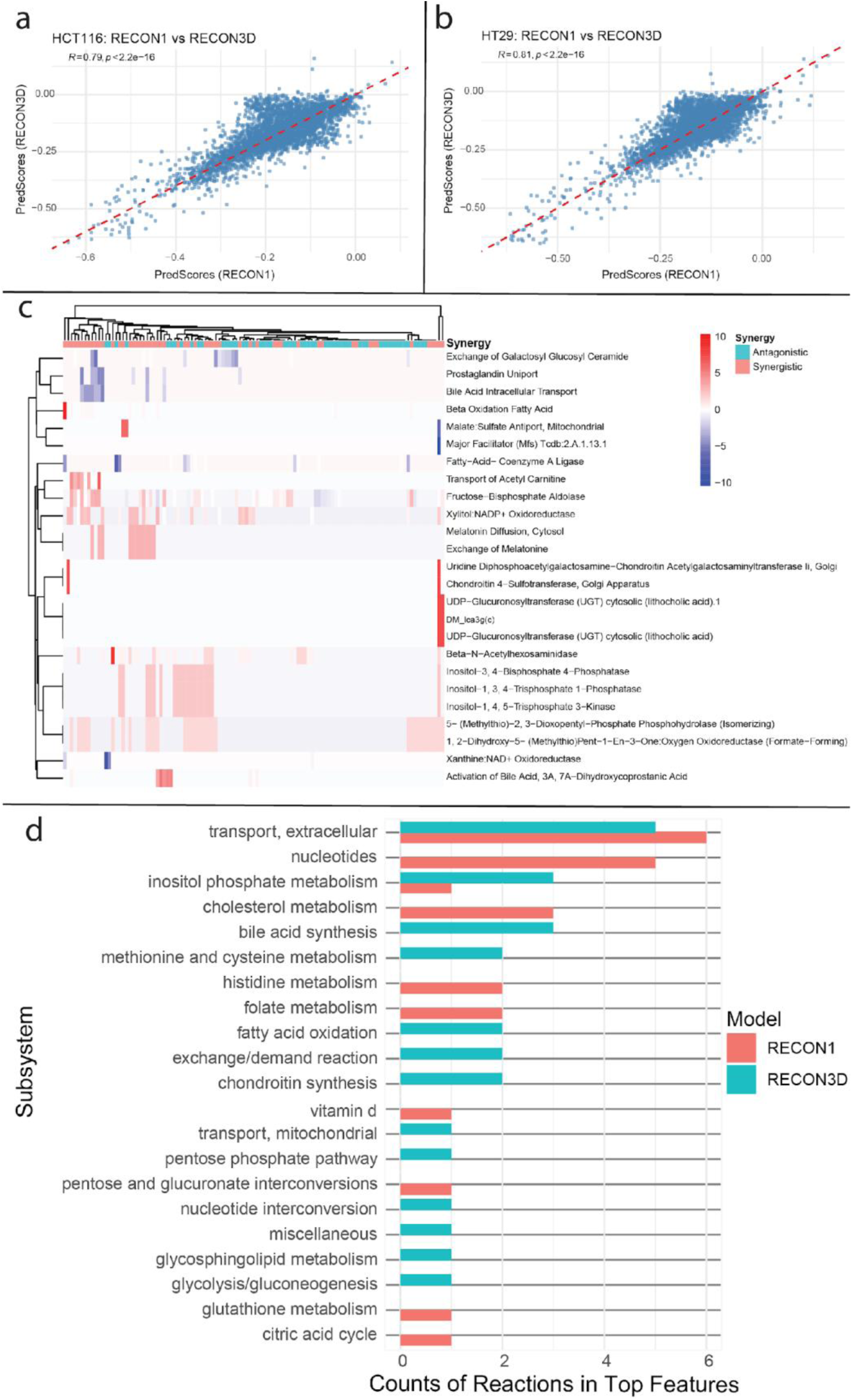
Comparison of RECON1 and RECON3D usage in OMG-ML predictions. (a-b) Comparison of predicted drug combination scores between models trained on RECON1 versus RECON3D metabolic networks. Each point represents a predicted combination score for a given cell line (HT29 or HCT116). The dashed diagonal line represents perfect agreement between the two sets of predictions. **(c)** Flux through the top 25 features for all drug treatments. Flux values were normalized before plotting. Synergy and antagonism classifications were made by averaging scores for all associated combinations for that drug. **(d)** Frequency of reaction subsystems in the top 25 features for the RECON1 and RECON3D trained models.

**Supplementary Fig. 4.**
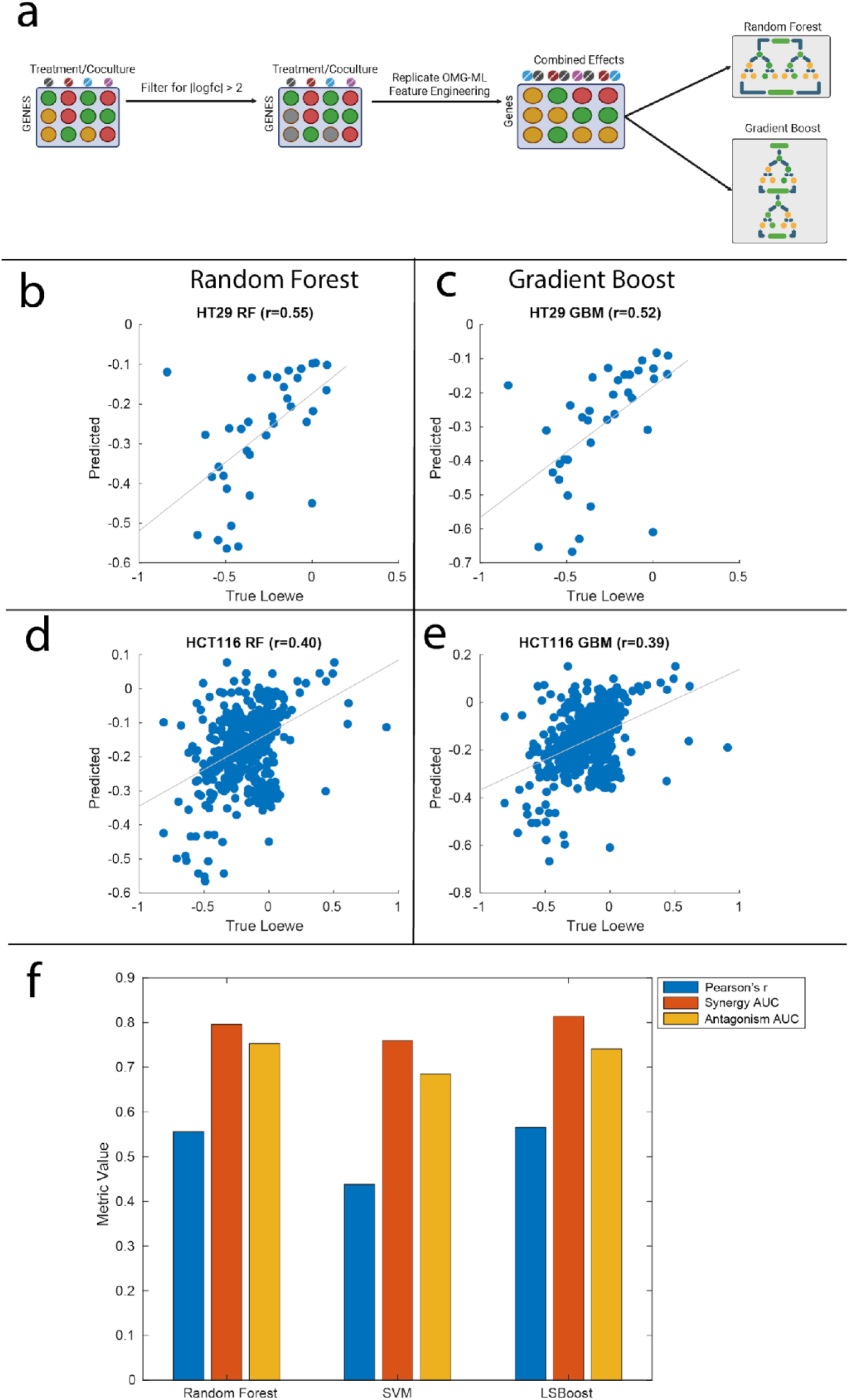
Methodology optimization. **(a)** Method for producing an ML model to predict drug synergy based on differentially expressed genes (without calculating flux). Feature engineering was performed to match OMG-ML feature engineering by simulating combined effects of drug combinations. These features were a sigma feature (characterizes combined effect of drugs), delta feature (characterizes how drug effects diverge), and an entropy feature (characterizes the diversity of gene perturbations by the drug treatment). Two separate model types were made, a random forest and a gradient boosted model. *Figure created using BioRender.com.* (**b-e)** Predicted scores by random forest and gradient boost model compared to true synergy scores from a withheld test set. Plots separated by cell type and ML type. **(f)** Compared performance of modeling types from 10 fold cross validation using OMG-ML model with flux input.

**Supplementary Fig. 5.**
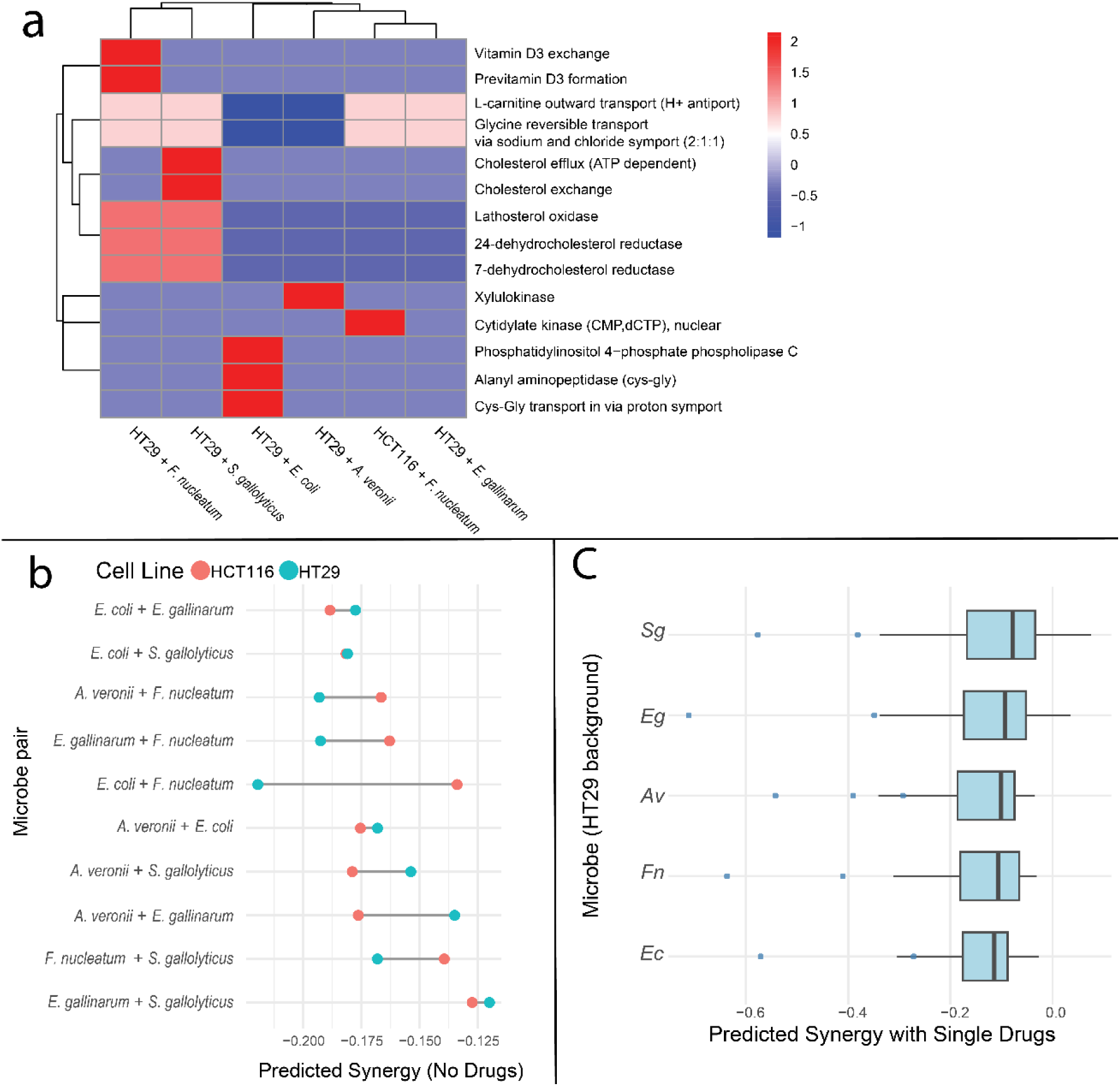
Metabolic effects and predicted synergy of individual microbes. **(a)** Z-score normalized flux through top features for co-culture-treated CRC cell lines. The top 25 features were filtered for the reactions where at least one treatment had flux. The drug with the most negative predicted score for all microbe treatments is cabazitaxel. **(b** Predicted synergy of two microbe combinations in the absence of drugs on HT29 cells compared to HCT116 cells. More negative scores indicate synergy. **(c)** Predicted synergy of 110 individual drugs with individual microbes on HT29 cells. More negative values indicate synergy. Of the co-cultured microbes, Fn had the most negative median score, meaning it had the most synergistic combinations (-0.20).

**Supplementary Table 1:** Top 15 most synergistic predicted drug combinations for HT29 and HCT116 cells.

| Top 15 Predicted Drug Synergies |  |  |  |
| --- | --- | --- | --- |
| HT29 and HCT116 Cell Lines |  |  |  |
| Drug 1 | Drug 2 | Predicted Synergy Score | Cell Line |
| cabazitaxel | erlotinib | -0.616 | HT29 |
| cabazitaxel | navitoclax | -0.608 | HT29 |
| cabazitaxel | pazopanib | -0.596 | HT29 |
| cabazitaxel | mk1775 | -0.595 | HT29 |
| cabazitaxel | estramustinephosphatesodium | -0.586 | HT29 |
| cabazitaxel | dabrafenib | -0.575 | HT29 |
| azd8055 | cabazitaxel | -0.571 | HT29 |
| cabazitaxel | methotrexate | -0.571 | HT29 |
| cabazitaxel | trametinib | -0.570 | HT29 |
| azd7762 | cabazitaxel | -0.565 | HT29 |
| cabazitaxel | taselisib | -0.560 | HT29 |
| cabazitaxel | mk2206 | -0.543 | HT29 |
| cabazitaxel | methoxsalen | -0.537 | HT29 |
| cabazitaxel | megestrol | -0.536 | HT29 |
| cabazitaxel | gemcitabine | -0.533 | HT29 |
| cabazitaxel | lenalidomide | -0.639 | HCT116 |
| cabazitaxel | erlotinib | -0.588 | HCT116 |
| cabazitaxel | navitoclax | -0.572 | HCT116 |
| cabazitaxel | trametinib | -0.556 | HCT116 |
| ixabepilone | thalidomide | -0.556 | HCT116 |
| cabazitaxel | dabrafenib | -0.547 | HCT116 |
| cabazitaxel | mk1775 | -0.542 | HCT116 |
| cabazitaxel | temozolomide | -0.541 | HCT116 |
| cabazitaxel | thalidomide | -0.537 | HCT116 |
| azd8055 | cabazitaxel | -0.533 | HCT116 |
| imiquimod | ixabepilone | -0.526 | HCT116 |
| cabazitaxel | taselisib | -0.525 | HCT116 |
| cabazitaxel | estramustinephosphatesodium | -0.524 | HCT116 |
| azd7762 | cabazitaxel | -0.520 | HCT116 |
| cabazitaxel | methotrexate | -0.518 | HCT116 |

